# The Vagus Nerve Mediates the Physiological but not Pharmacological Effects of PYY_3-36_ on Food Intake

**DOI:** 10.1101/2020.08.07.241851

**Authors:** A Martin Alonso, SC Cork, Y Ma, M Arnold, H Herzog, SR Bloom, W Distaso, KG Murphy, V Salem

## Abstract

Peptide YY (PYY_3-36_) is a post-prandially released gut hormone with potent appetite-reducing activity mediated by the neuropeptide Y (NPY) Y2 receptor (Y2R). However, the neuronal pathways by which PYY_3-36_ acts to supress appetite are unclear. Determining how the PYY_3-36_ system physiologically regulates food intake may help exploit its therapeutic potential. Here we demonstrate that germline and post-natal targeted knockdown of the Y2R in the afferent vagus nerve inhibits the anorectic effects of physiologically-released PYY_3-36_, but not peripherally-administered higher doses. Post-natal knockdown of the Y2R results in a transient body weight phenotype that is compensated for in the germline model. Loss of vagal Y2R signalling also alters meal patterning and accelerates gastric emptying. These results may facilitate the design of PYY-based anti-obesity agents.

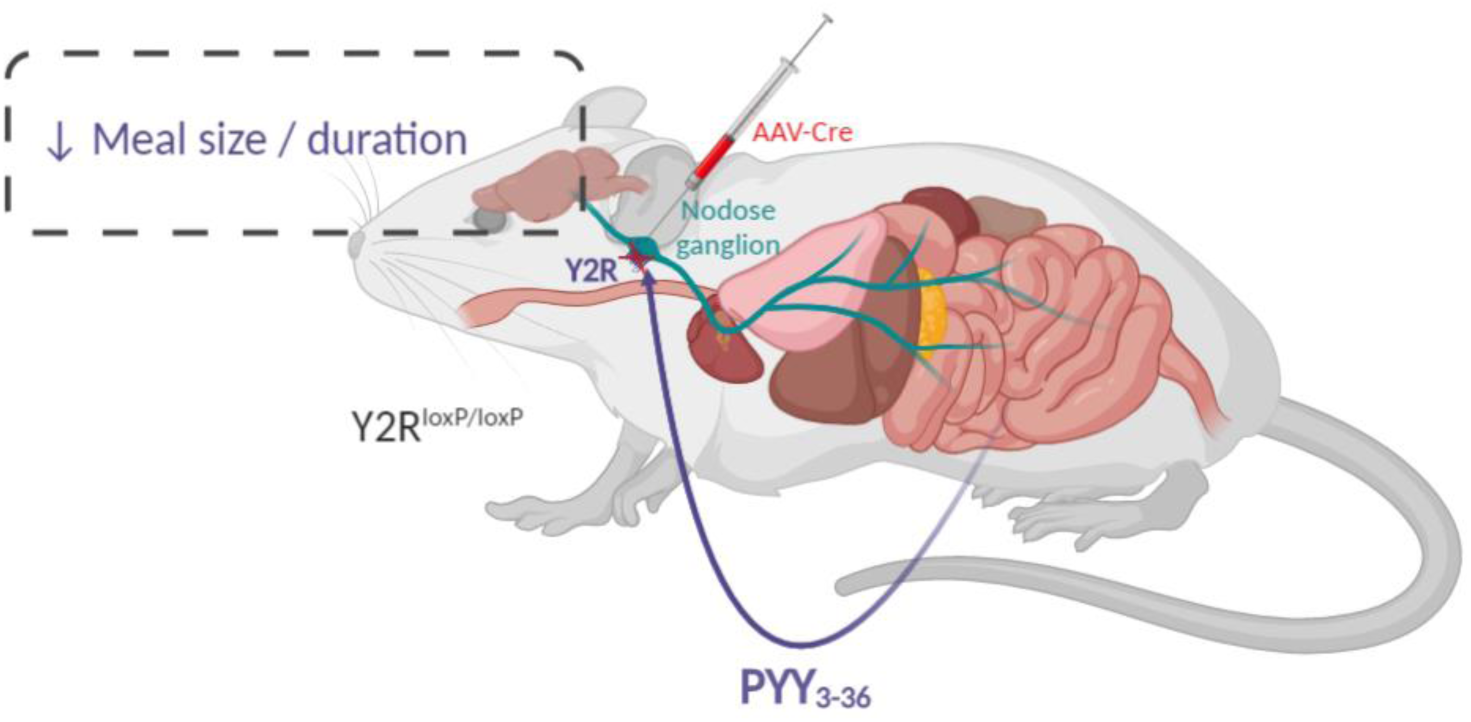

---

The anorectic gut hormone peptide YY (PYY) is secreted from enteroendocrine L-cells (EECs) in response to food intake in a manner dependent on both caloric load and macronutrient composition^1,2,3^. Peripheral administration of the active circulating form, PYY_3-36_, powerfully reduces food intake in rodents and humans, and chronic administration induces weight loss in rats^4,5^. PYY_3-36_ retains its anorectic effects in obese subjects^6^, suggesting possible utility as an anti-obesity pharmacotherapy. However, the mechanism of action of this peptide is not fully understood. Given the challenges of maintaining the effects of PYY_3-36_ chronically, and potential side effects including nausea, understanding how this system physiologically regulates food intake may help unlock its therapeutic potential whilst minimising unwanted effects.

PYY_3-36_ mediates its anorectic effects via the neuropeptide Y (NPY) Y2 receptor (Y2R)^4^, but the site of action is unclear. Pharmacological administration studies have reported direct effects of PYY_3-36_ on the brain^4,7^. It has been postulated that circulating PYY_3-36_ crosses the blood-brain barrier and directly accesses the Y2R in the hypothalamic arcuate nucleus (ARC) to inhibit feeding^4,7,8^. Here, Y2R acts as an auto-inhibitory presynaptic receptor in arcuate NPY neurons. PYY therefore inhibits NPY neurons resulting in the loss of tonic inhibition of neighbouring anorexigenic proopiomelanocortin (POMC) neurons^4^.

However, a number of studies suggest that PYY_3-36_ does not act directly on the hypothalamus to suppress food intake^9,10^. PYY_3-36_ retains its anorectic effect in POMC-deficient and melanocortin-4 receptor (MC4R)-deficient mice^11,12^, suggesting that the melanocortin system is not essential for PYY_3- 36_-induced suppression of food intake.

The vagus nerve, the main neural link between the gut and the brain, also expresses Y2R in rodents^9,13^ and in humans^13^. Furthermore, Y2R expression levels in the vagus are regulated by nutritional status^13^.

The necessity for intact vagal signalling for the anorectic effects of peripheral PYY_3-36_ is unclear, with studies reporting an abrogation of its effects following subdiaphragmatic vagotomy in rats^9,14^ but not mice^10,15^. These studies all used vagal lesioning methods, which prevent all vagal afferent and efferent signalling, and can therefore result in confounding effects on appetite and gastrointestinal function that make it difficult to determine whether PYY_3-36_ mediates its physiological effects on food intake via the vagus.

We hypothesised that the physiological anorectic effects of PYY_3-36_ are mediated by the vagus nerve. Here we demonstrate that animals with germline knockdown of the NPY Y2R in sensory neurons (Nav1.8/Y2R KO), including the afferent vagus, demonstrate an attenuated response to low but not high dose PYY_3-36_. To avoid the possible confounding effects of developmental compensation, we then generated a mouse model of adult Y2R knockdown specific to the afferent vagus. We used bilateral injection of an adeno-associated virus expressing cre recombinase (AAV-Cre) into the nodose ganglia (NG), extracranial structures that contain the cell bodies of vagal afferents, to achieve selective knockdown of the Y2R in the vagal sensory neurones of mice with a floxed Y2R (NG Y2R KD). We demonstrate that postprandially-released PYY_3-36_ requires intact vagal signalling but that pharmacological doses suppress appetite despite loss of vagal afferent Y2R.

Nav1.8 is a sodium channel isoform expressed in sensory neurons of the vagus nerve (over 75% of mouse vagal neurones) as well as in the spinal cord^16^. Y2R^loxP^ mice were bred with mice expressing cre recombinase driven by the Nav1.8 promoter to generate Nav1.8/Y2R KO animals. In this germline model, Y2R mRNA expression was significantly reduced in both the left and right NG of Nav1.8/Y2R KO compared to littermate controls (**Fig. 1a**). No differences in adult body weight or body composition were noted between Nav1.8/Y2R KO and control mice (**Fig 1b**).

**Figure 1.**
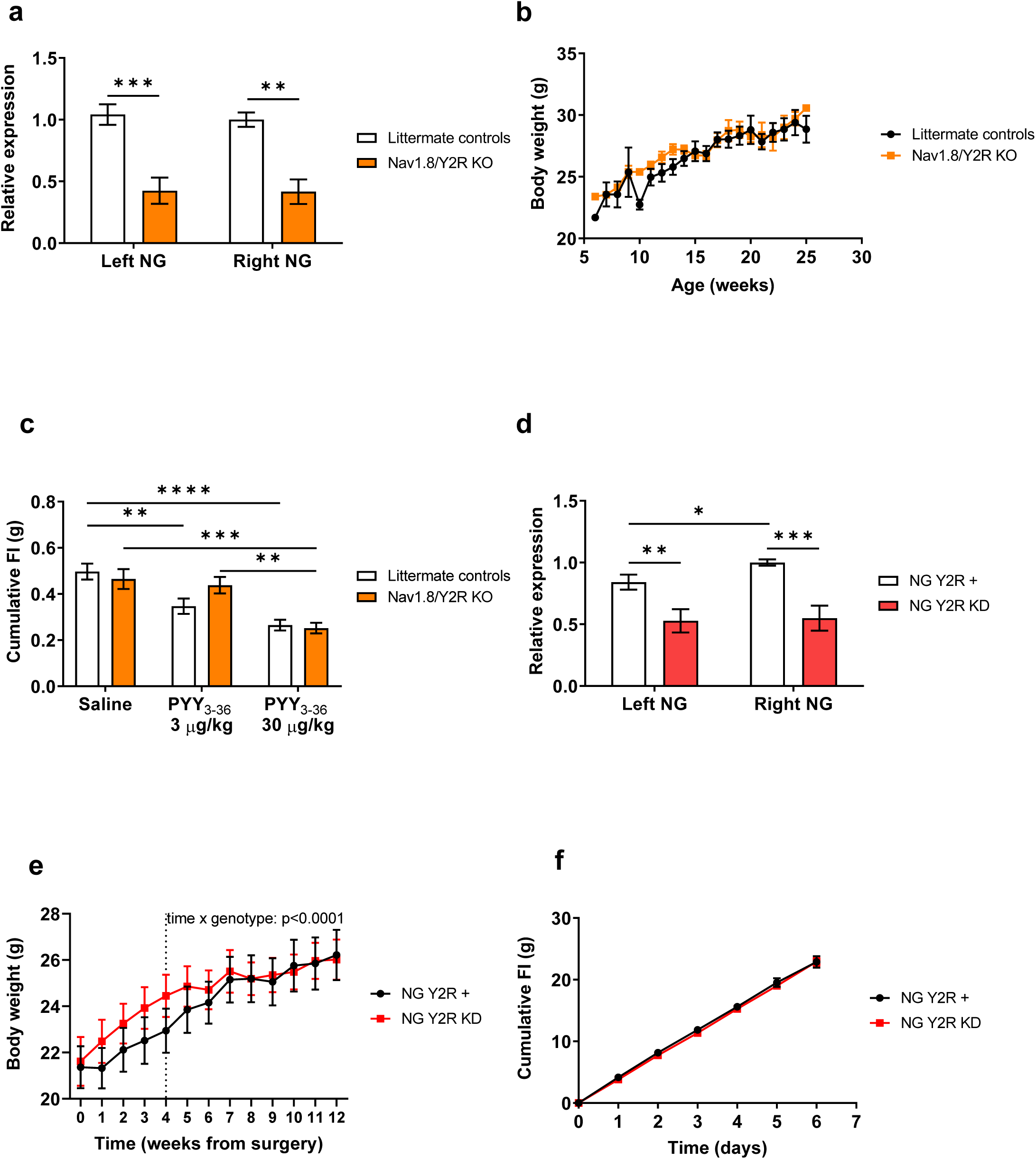
**1a.** Relative expression of Y2R mRNA in left and right NG of Nav1.8Cre/Y2R^loxP/loxP^ mice (Nav1.8/Y2R KO) and littermate controls that were sacrificed at 16-30 weeks of age (n = 4-5 mice). Expression was normalised to the Y2R expression in the right nodose ganglion (NG) of control mice. **p < 0.01, ***p < 0.001 by two-way ANOVA followed by Bonferroni’s test. **1b**. Body weight at 5-25 weeks of age in Nav1.8/Y2R KO and littermate control mice (n = 7-12 mice). **1c**. One-hour food intake (FI) after vehicle or PYY_3-36_ (3 μg/kg or 30 μg/kg, IP) administration in the early dark-phase in Nav1.8/Y2R KO vs littermate control (n = 7-11). **p < 0.01, ***p < 0.001, ****p < 0.0001 by two-way ANOVA followed by Bonferroni’s test. **1d**. Relative expression of Y2R mRNA in left and right NG of Y2R^loxP/loxP^ mice injected with AAV-Cre into the NG (NG Y2R KD) and of littermates injected with control virus (AAV-GFP) into the NG (NG Y2R +) that were sacrificed at 17-21 weeks post-surgery (n = 13-14 mice). Expression was normalised to the Y2R expression in the right NG of control mice. *p < 0.05 by two-tailed Student t test. **p < 0.01, ***p < 0.001 by two-way ANOVA followed by Bonferroni’s test **1e**. Body weight post-surgery in NG Y2R + and NG Y2R KD mice 0-12 weeks post-surgery (n = 17-15 mice). p < 0.0001 by two-way ANOVA. **1f**. Food intake in NG Y2R + and NG Y2R KD mice over 6 days at week 7 post-surgery.

We investigated whether the response to exogenous PYY_3-36_ was dependent on vagal Y2R signalling. The effects of intraperitoneal administration of low and high dose PYY_3-36_ was investigated in Nav1.8/Y2R KO animals compared with littermate controls. The low dose (3 µg/kg) had previously been shown not to cause conditioned taste aversion ^10^, and was compared to a high dose (30 µg/kg) more likely to access centrally expressed Y2R^4,10,12^.

Interestingly, low dose PYY_3-36_ suppressed food intake in littermate controls but not in Nav1.8/Y2R KO mice. In contrast, high dose PYY_3-36_ suppressed food intake in both groups **(Fig 1c)**. These results suggest that low-dose exogenous PYY_3-36_, which may better reflect the actions of endogenous PYY_3-36_, acts via Y2R vagal pathways, but that high-dose PYY_3-36_ bypasses vagal signalling.

To confirm that these findings did not reflect developmental differences, we generated a vagus-specific, adult knockdown model by administering bilateral intra-NG injections of AAV-Cre in Y2R^loxP/loxP^ mice on a C57/Bl6J background (NG Y2R KD). This resulted in a significant bilateral reduction in Y2R mRNA in the NG, as measured at 20-24 weeks post injection, compared to controls injected with adeno-associated virus expressing green fluorescent protein, AAV-GFP (NG Y2R +) (**Fig. 1d**). Animals were group-housed in the early postoperative period for welfare reasons. Body weight was significantly higher in the NG Y2R KD group up to 6 weeks following injection, when the body weight curves converged (**Fig 1e**). By week 7 post injection, body weight and food intake in each group (NG Y2R KD vs NG Y2R +) was similar (**Fig 1f**).

To confirm that the Y2R knockdown was specific and that other vagal afferent signalling pathways remained intact following NG injection, we investigated the response to cholecystokinin octapeptide (CCK-8), a gut peptide established to exert anorectic effects via vagal afferents^17^. CCK-8 at 5 µg/kg (intraperitoneal, IP) significantly reduced subsequent food intake compared to vehicle in both groups (p = 0.0044) and no difference (p = 0.4657) was observed in CCK-8-mediated food intake reduction between NG Y2R KD and NG Y2R + mice (**Fig 2a**).

**Figure 2:**
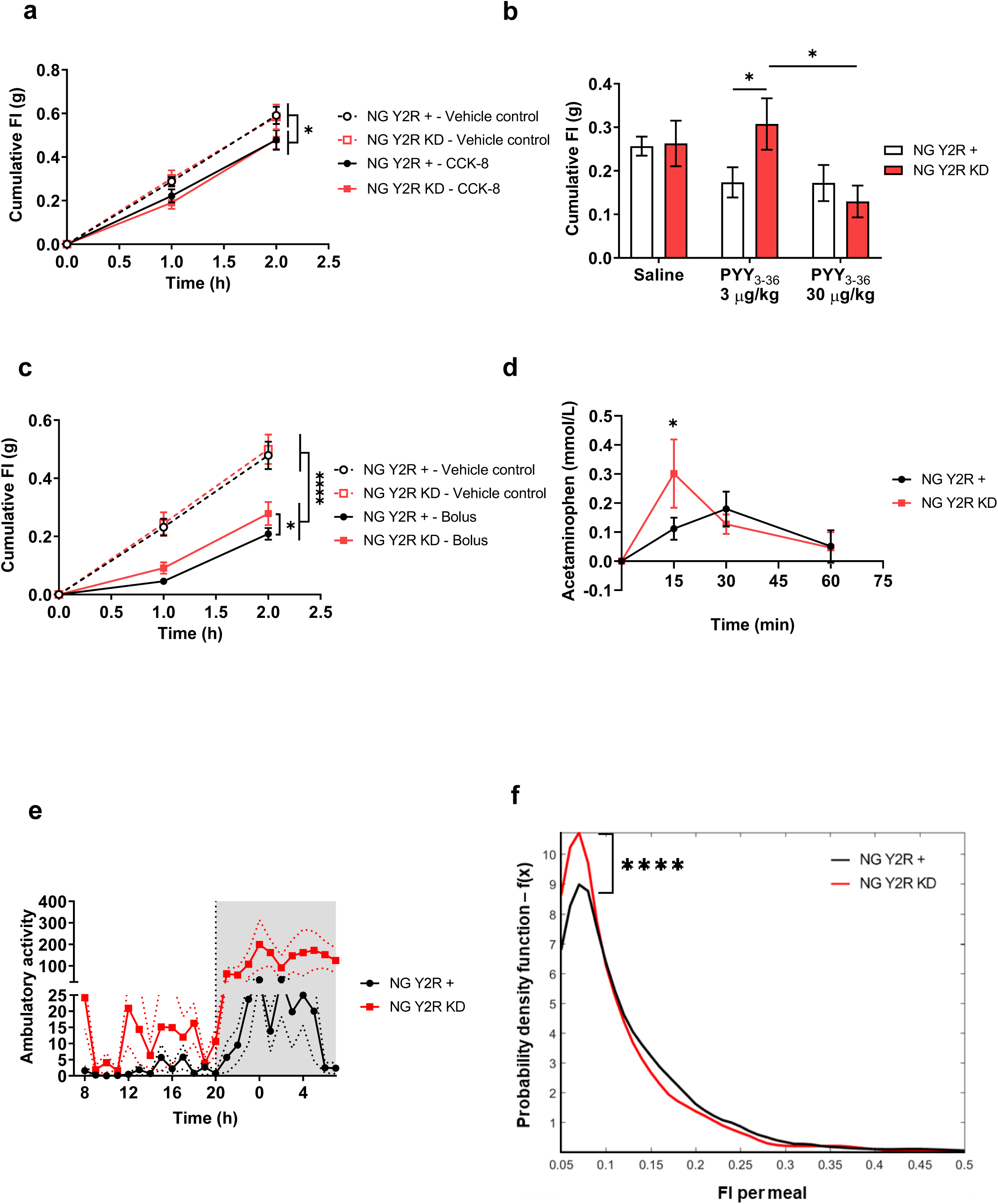
**2a.** One-hour food intake (FI) after vehicle or CCK-8 (5 μg/kg, IP) administration in the early dark phase (n = 13-14). *p < 0.05 (time x treatment) by three-way ANOVA. Two-way ANOVA followed by Dunnett’s test to compare both time 1h and 2 h to time 0 h resulted in p < 0.0001 for both time points in both control and NG Y2R KD. **2b**. One-hour FI after vehicle or PYY_3-36_ (3 μg/kg or 30 μg/kg, IP) administration in the early dark-phase in (AAV-Cre injected) NG Y2R KD vs littermate control animals (n = 7-9). *p < 0.05 by two-way ANOVA followed by Bonferroni’s test or by multiple t test with Bonferroni’s test. **2c**. The feeding response to a gavage of either a high calorie meal or an equal volume of water was compared in NG Y2R KD and NG Y2R + control (AAV-GFP injected) mice. Cumulative food intake (FI) over 2.5 hours after gavage of water is shown in on the dashed lines, with no differences observed between the NG Y2R KD (orange) and control groups. However, after gavage of a caloric meal, cumulative 2.5 hour FI in the early dark phase was significantly greater in the NG Y2R KD group in line with the hypothesis that vagal Y2R mediates the anorectic effects of postprandially release PYY (n = 13-12). ****p < 0.0001 (time x treatment) by three-way ANOVA. *p < 0.05, **p < 0.01 by two-way ANOVA followed by Bonferroni’s test. Two-way ANOVA followed by Dunnett’s test to compare to time 0 (at t = 1h in Bolus condition, p < 0.01 for both groups; at t = 1h in the control condition, p < 0.001 for NG Y2R + and p < 0.0001 for NG Y2R KD; at t = 2h, p < 0.0001 in all conditions and groups). **2d**. Plasma acetaminophen levels after oral gavage of a 20% glucose bolus containing 1% acetaminophen in 9-h fasted animals (n = 6-3). *p < 0.05 by multiple t test with Bonferroni’s test. **2e**. Ambulatory activity (beam breaks) in NG Y2R KD and controls (mean of 3 days, dark phase in grey) in the CLAMS system **2f**. Probability density function of food intake per meal over 7 days in a CLAMS system measuring high resolution food intake. The difference between NG Y2R KD vs control animals is highly significant (p<0.0001) Kolmogorov-Smirnov (K-S) test (n = 6 and 6).

The dose-dependent effects of PYY_3-36_ were recapitulated in this more specific vagal Y2R knockdown model. In control animals, peripheral injection of low-dose PYY_3-36_ tended to reduce food intake over the next hour (p = 0.059, n = 9) compared to vehicle, but this effect was significantly abrogated in NG Y2R KD mice. However, high-dose PYY_3-36_ produced a similar reduction in food intake in both NG Y2R KD (−0.133±0.064 g, n = 7, p = 0.058) and control-injected NG Y2R + animals (−0.085±0.047 g, n = 9, p = 0.089) compared to saline (**Fig 2b**).

To study the role of the vagal Y2R in response to physiological PYY release, postprandial food intake was measured following oral gavage of a mixed nutrient meal (5.3 kcal/mL, energy: 22% CHO, 69% Fat, 9% Protein). This meal was confirmed to induce a more than six-fold rise in total plasma PYY levels in wild-type C57 mice, from 10.10±9.21 pmol/L to 69.95±21.60 pmol/L at 30 minutes post gavage (n = 4-5; p = 0.0534) **(Supp Fig. 1a)**. This rise in PYY is similar to that previously reported in mice and in humans postprandially^1^. A*d libitum* food intake was significantly reduced at 2 hours compared to gavage of vehicle control in both NGY2R + (−0.267±0.051 g, p <0.0001, n = 13) and NG Y2R KD mice (− 0.221±0.064 g, p = 0.002, n = 12), in accord with mechanical distension and the release of other anorectic gut hormones besides PYY reducing appetite. However, this suppression of food intake was significantly attenuated in the NG Y2R KD compared to NG Y2R + mice (p = 0.044) (**Fig 2c**). Moreover, the percentage knockdown of Y2R achieved in the NG significantly correlated with the degree of attenuation of food intake suppression (R=0.4, p=0.04). These data strongly suggest that vagal Y2R signalling plays a physiological role in post-prandial satiation.

Previous studies have suggested that PYY_3-36_ inhibits gastric emptying^18^, but the pathway mediating this effect is unknown. To investigate whether delayed gastric emptying might play a role in vagal Y2R-mediated feeding suppression, we performed an acetaminophen absorption test in a separate cohort of NG Y2R KD and control-injected NG Y2R + mice. This revealed a significantly higher peak of blood acetaminophen at 15 minutes in the NG Y2R KD group compared to control injected animals, suggesting quicker gastric emptying (**Fig 2d**). A trend towards an increased area under the curve (AUC) for glucose (278.4 vs 409.9 mmol/L x time; p = 0.1275) in an oral glucose tolerance test (OGTT) was also observed in the NG Y2R KD group **(Supp Figs 1b and 1c**). Together, these results suggest that PYY_3-36_ released in response to nutrients in the gut slows gastric emptying by acting on vagal afferents, an effect that may contribute to its anorectic effects.

Finally, we investigated whether the vagal Y2R modulated meal patterning rather than overall energy intake. Control-injected NG Y2R + and NG Y2R KD animals were studied in a Comprehensive Laboratory Animal Monitoring Systems (CLAMS) system for 10 days, 10-12 weeks after successful AAV-cre injection. A trend towards increased ambulatory activity was observed in the dark phase in NG Y2R KD (**Fig 2e and Suppl Fig 1d**). No differences in overall food intake, energy expenditure (EE), respiratory exchange ratio (RER) or oxygen consumption were observed in NG Y2R KD animals compared to NG Y2R + controls. However, probability density functions for meal patterning readouts revealed highly significant differences (p <0.0001) in average meal size and duration; meals were smaller and shorter in NG Y2R KD compared with controls (**Fig. 2f and Suppl Fig 1e**), with shorter intermeal durations. This was coupled with a non-significant tendency (p = 0.09) for more meals in the KD group over the duration of the CLAMS experiment. These data suggest a physiological role for vagal Y2R on meal patterning, even in the absence of an overall long-term effect on energy balance.

The contribution of gut hormones in the regulation of short-term food intake and longer-term energy homeostasis is well established, and the potential for targeting these hormone systems to promote weight loss has been realised with the licencing of GLP-1 analogues to treat obesity^19^. However, the pathways involved in gut hormone signalling, and the physiological effects of individual hormones are unclear. In particular, the complex role of the vagus nerve in the gut-brain axis is just beginning to be understood. Only recently have new techniques in neuroscience and genetics successfully profiled vagal innervation of the gut and established the function of individual neuronal populations^20,21^. Enteroendocrine cells in the gastrointestinal tract postprandially secrete satiety-inducing gut hormones, the receptors for which are expressed in vagal afferents^22^. Gut hormones may diffuse through the lamina propria and interact with vagal afferents terminals, or neuropods on enteroendocrine cells themselves may directly synapse with the vagus nerve^23^.

PYY and GLP-1 are co-secreted from L-type EECs and both modulate food intake in humans^24,25^. Our results add to the body of literature suggesting that individual gut hormones act, at least in part, through the vagus nerve to modulate feeding behaviours^26,27^. A recent study suggests that PYY, and not GLP-1, from colonic EECs is responsible for the appetite suppression observed following the activation of these cells^28^. The current study suggests that physiologically released PYY_3-36_ requires intact vagal signalling to mediate its anorectic effects, but that exogenous administration of higher, pharmacological doses bypasses this pathway.

There is evidence that gut hormones acting physiologically via the vagus nerve may modulate the sensitivity of mechanosensitive vagal afferent endings^26,27^. Here we demonstrate that Y2 receptor depletion in the vagus nerve modulates meal patterning in association with accelerated gastric emptying. Animals with knocked down Y2R in the vagus engaged in shorter and quicker meals, with reduced times between feeding bouts, suggesting that their gastric emptying was accelerated.

Mice congenitally lacking the Y2R in their sensory afferent nerves as well as those with adult knockdown of the receptor demonstrated an abrogated response to low dose PYY but a preserved response to high dose PYY. Congenital Nav1.8 driven loss of Y2R in vagal afferents did not produce an adult body weight or cumulative food intake phenotype compared with littermate controls. This may have been because of the incomplete knockdown of receptors and some redundancy in the system or because of compensatory sequelae. However, mirroring the transient effects of virally-induced knockdown of the hypothalamic Y2R^29^, we observed a body-weight phenotype that suggested a positive energy balance in the vagal Y2R KD animals for the first 6 weeks of cre expression, in a post natal model that achieved similar knockdown efficiency. Since individual food intake was not measured during this period of group housing, it remains unclear whether this was due to an associated increase in food intake, as might be expected in a model that blocks the signalling of an anorectic hormone. By 12 weeks post injection, the body weight and food intake phenotype of the vagal Y2R KD animals had reverted to that of the control treated animals, though there was still a clear difference in meal patterning behaviour and gastric emptying. The early and short-lived body weight phenotype elicited in the post-natal knockdown model supports the notion of compensatory mechanisms with single receptor knock down models, particularly in complex systems such as energy homeostasis. Further understanding of the phenotype and distribution in vagal afferent endings within the gut would help to distinguish the relative advantages of the Nav1.8 versus the NG injection models. Nav1.8-positive neurons are observed in hormone-sensing mucosal endings as well as in a subgroup of tension-sensing vagal intraganglionic laminar endings, and their anatomical proximity to enteroendocrine cells may vary by neuronal type^30^. It is also noteworthy that the low degree of knockdown achieved in both Y2R KD models might suggest that a more severe knockdown of vagal Y2R might result in a more characteristic phenotype and that the role of PYY signalling in the vagus nerve might be greater.

In summary, these data suggest that Y2R signalling in the vagus nerve plays a physiological role in mediating the anorectic and meal patterning effects of endogenous PYY_3-36_.

## Acknowledgments

The Section of Endocrinology and Investigative Medicine is funded by grants from the MRC, BBSRC, NIHR and is supported by the NIHR Biomedical Research Centre Funding Scheme. The views expressed are those of the authors and not necessarily those of the MRC, BBSRC, the NHS, the NIHR or the Department of Health and Social Care. SCC is supported by a project support grant by the British Society for Neuroendocrinology and a Wellcome Trust Institutional Strategic Support Fellowship. KGM is supported by a Diabetes UK project grant. VS is funded by a Harry Keen Diabetes UK Fellowship.

## Author Contributions

Conceptualization VS; Methodology, AMA, SC, YM, MA and VS; Formal analysis, AMA, SC, WD, VS; Data curation VS; Writing—original draft preparation, AMA, SC, KGM and VS; Writing—review and editing, all authors; Visualization, WD and VS. All authors have read and agreed to the published version of the manuscript.

## Methods

### Animals

To establish the Y2R floxed mice colony, sperm from Y2R^loxP/loxP^ mice (generated by Prof Herbert Herzog, Garvan Institute of Medical Research) was used to fertilize C57BL/6J oocytes. Littermates Y2R^loxP/loxP^ and Y2R^loxP/-^ were used as controls^1^.

To selectively delete Y2R in primary afferent neurons from birth, the Nav1.8/Y2R KO mouse line was generated. Nav1.8 knock-in Cre-recombinase mice on a C57Bl/6J background were previously generated by Prof John Wood (University College London)^2^. C57BL/6J oocytes were fertilised with sperm from Nav1.8^Cre^ mice (MRC Harwell Institute, Oxfordshire, UK). Mice were bred to generate Nav1.8^Cre^ hemizygous Y2R^loxP/loxP^. Littermates Cre-negative Y2R^loxP/loxP^, Nav1.8^Cre^, and Nav1.8^Cre^ Y2R^loxP/-^ were used as controls in the studies.

All mice were housed under controlled temperature (21-23°C) and a 12-h light/dark cycle schedule (lights on 7:00 - 19:00 or 23:00 - 11:00) with *ad libitum* access to water and standard chow RM1 (SDS Diets, Witham, UK) unless specified otherwise. All animal procedures complied with the British Home Office Animals (Scientific Procedures) Act 1986 regulations and were approved by the Animal Welfare and Ethics Review Board at Imperial College London.

### Nodose ganglia microinjection

Adult mice (6-14 weeks old, 22.2±0.6 g on surgery day) were anaesthetised with 5% isoflurane (IsoFlo®, Zoetis, London, UK) and 1L/min oxygen in an anaesthetic chamber. Mice received subcutaneous (s.c.) injections of 2 mg/kg atropine sulfate (Hameln Pharmaceuticals, Gloucester, UK) and 5 mg/kg analgesic carprofen (Rimadyl®, Zoetis) at the onset of surgery. Mice were placed in a supine position and anaesthesia was maintained by mask inhalation of isoflurane. The NG was exposed so it was accessible for injection. A previously pulled 0.58 mm-diameter glass capillary (World Precision Instruments, Hertfordshire, UK) containing AAV was attached to a micromanipulator and the NG was injected (5nL) using a PV820 Pneumatic Picopump (World Precision Instruments). After the surgery, mice received s.c. injection of 0.05 mg/kg buprenorphine hydrochloride (Vetergesic®, Ceva Animal Health, Buckinghamshire, UK). Mice recovered in a ventilated heating chamber and were returned to their group home cage. Animals body weight and state were daily recorded until BW values reached pre-surgery levels. After a recovery week, surgery was repeated on the other NG.

Animals recovered for four weeks to allow expression of the viral constructs. The viruses used were AAV9.hSyn.eGFP.WPRE.bGH (titre: 1.32×10^14^ GC/mL), AAV9.hSyn.HI.eGFP-Cre.WPRE.SV40 (titre: 5.506 ×10^13^ GC/mL), AAV5.hSyn.Cre.hGH (titre: 2.484×10^13^ GC/mL) and AAV5.hSyn.eGFP.WPRE.bGH (titre: 2.224×10^13^ GC/mL). All the viruses were supplied by Penn Vector Core (Gene Therapy Program, University of Pennsylvania, US).

### Gene expression analysis

Mice were genotyped using the KAPA2G Fast HotStart Mouse Genotyping Kits (Merck, Southampton, UK) in a PCR. Y2R floxed genotype was confirmed using primer 5′-TTAACATCAGCTGGCCTAGC-3′, and 5′-GGAAGTCACCAACTAGAATGG-3′^3^. Nav1.8-Cre were genotyped using the following primers: a) 5′-CAGTGGTCAGGCTGTCACCA-3′ (common forward); b) 5′-ACAGGCCTTCAAGTCCAACTG-3’ (reverse wild-type); c) 5′-AAATGTTGCTGGATAGTTTTTACTGCC-3′ (reverse Cre). All primers were supplied by Sigma A-ldrich (Dorset, UK).

Tissue was collected after IP overdose of anaesthetic. Both NG were collected and snap frozen. RNA was extracted after tissue homogenisation in TRIsure™ (Bioline, London, UK) using a TissueLyser II (Qiagen, Manchester, UK). DNA was digested using RQ1 RNase-Free DNase kit (Promega, Hampshire, UK). High Capacity cDNA Reverse Transcription Kit (Applied Biosystems, Warrington, UK) was used to synthetize cDNA from 100 ng of RNA. Quantitative PCR (qPCR) was performed in triplicates using TaqMan assays (Applied Biosystems) in a CFX384™ detection system (Bio-Rad, Hertfordshire, UK). Taqman probes used were neuropeptide Y receptor Y2 (Npy2r; Mm01956783_s1) and peptidylprolyl isomerase (cyclophilin)-like 3 (Ppil3; Mm00510343_m1). Relative levels of mRNA gene expression were calculated using the 2^−ΔΔCt^ method.

### Feeding studies

In order to acclimatise the animals, mice were singly caged and accustomed to IP injection (by injections of 100 μL of saline) or oral gavage (by oral gavaging of 300 μL of water) in the early dark phase every other day during the week before the studies.

Using a crossover design, animals received each treatment in a random order. In the case of feeding studies following intraperitoneal (IP) administration, the treatments used were PYY_3-36_ (Tocris Biosciences, Abingdon, UK), sulfated cholecystokinin octapeptide CCK-8 (CCK-8; Tocris Bioscience) or saline (vehicle control). Treatments were prepared by adding saline to freeze-dried vials or directly to the lyophilized peptide. Mice were IP injected with a volume of 100 μL. In the case of feeding studies following oral gavage, mice received 300 μL gavage of either the liquid mixed nutrient meal or control (water). The gavaged meal composition was 17% (w/v) of D(+)-Glucose (Sigma-Aldrich), 8.3% (w/v) of L-arginine (Sigma-Aldrich), 41.7% (v/v) of extra virgin olive oil (Sainsbury’s Supermarkets Ltd, London, UK) and 58.3% (v/v) in Ensure® Plus (Abbott, Maidenhead, UK).

Treatments were prepared immediately prior to the study. Mice were fed *ad libitum* both before and during the study. Just before the onset of the dark phase, animals were weighed and food (including spillage) was removed so no food was available in the cages until the treatment. In the early dark phase (t = 0 h), anorectic drugs or bolus were administered. Immediately thereafter, pre-weighed standard chow RM1 was placed on the hopper feeder and the food content in each cage was weighed using a PLJ 420-3F (Kern & Sohn GmbH, Germany) precision balance (0.001 g readability). Food intake was measured as the difference between the initial chow weight and chow weight (plus spill) remaining at the different time points post-treatment. For chronic cumulative food intake recording, chow (RM1) was weighed daily at the same time of the day using the same precision balance as for acute feeding studies.

### Indirect calorimetry and assessment of physical activity

NG Y2R KD and NG injected controls (NG Y2r +) were housed individually at 19-27 weeks of age (>12 weeks after surgery) for 10 days in a Comprehensive Lab Animal Monitoring System (CLAMS, Columbus US). Body weight was measured at the beginning of the experiment (25.9±0.7 g). Mice were housed at 22°C with airflow of 0.5 l/min. Chow (RM1) and water were available *ad libitum*. Oxygen consumption rate (VO2) and carbon dioxide output were measured using an open-circuit indirect calorimeter (Oxymax series; Columbus Instruments) were measured every 27 min. Energy expenditure and respiratory exchange ratio (RER) was calculated. Locomotion was recorded continuously at the same time as the indirect calorimetry measurements. Total counts (every time a beam is broken) and ambulatory counts (when a consecutive adjacent beam is broken and, therefore, actual physical activity) in the x- and y-axes directions are recorded. High resolution food intake was possible on an integrated continuous food weight measurement system.

### PYY radioimmunoassay

In the early dark phase, mice were orally gavaged with 300 µl of a caloric bolus (see above). Blood samples were collected at 30 min and 1 h post-gavage in 500 µL Microvette® tubes containing anticoagulant K3 EDTA (Sarsted, Leicester, UK) and 10 µL (1,000 KIU) of the protease inhibitor Aprotinin (Nordic Pharma, Reading, UK). Plasma was isolated by centrifugation at 3200*xg* for 15 min and stored at -80°C.

Plasma PYY was measured using a previously established in-house radioimmunoassay^4^. The polyclonal anti-PYY antibody has 100% cross-reactivity with PYY_1-36_ and PYY_3-36_. The separation method was based on dextran-coated charcoal and assays were counted in the gamma counter Multi Crystal LB 2111 (Berthold Technologies, Harpenden, UK). The intra-assay coefficient of variation was 3.47%.

### Oral glucose tolerance test

Following 8 h-fasting with *ad libitum* access to water during the light phase, local anaesthetic cream was applied on the tail of mice and the baseline blood glucose level was measured following venesection and sampling from the tail tip using a glucometer (GlucoRx, Surrey, UK) 15 min before glucose (Sigma-Aldrich) administration. Then, 2 g/kg of D(+)-glucose (20% w/v) was administered by oral gavage and blood glucose levels were measured at 15, 30, 60, 90 and 120 min.

### Gastric Emptying

Following an 8 h-fasting in the light phase with *ad libitum* access to water (33-39 weeks of age), local anaesthetic cream was applied on the tail surface. At t = 0, blood from the tail vein was collected and animals received an oral gavage of a bolus containing 1% paracetamol (100 mg/kg) 20% D(+)-glucose. Blood from the tail vein was collected at 15, 30 and 60 min in Microvettes® CB 300 containing EDTA (Sarstedt Inc), spin at 3200*xg* for 15 min and plasma was stored at -80°C. The acetaminophen assay (Cambridge Life Sciences Ltd, Cambridgeshire, UK) was performed using 5 µL of plasma following a recommended reduced volume protocol.

### Statistics

Data from IP feeding studies are expressed as mean ± SEM. Statistical analysis was performed by two-tailed unpaired t-test or by one-way or two-way ANOVA after confirmation of normality (Pearson D’Agostino). When ANOVA indicated a significant difference in the main effect or interaction, groups were compared by Dunnett’s, Tukey’s or Bonferroni’s post hoc test. p<0.05 was considered significant. Graphs were generated using GraphPad Prism (version 8).

Data for high resolution food intake between the NG Y2R KD and control injected groups were first assessed for measurement errors. Using maximum cut offs defined by the Palmiter group for normal meal sizes and duration in mice^5^, less than 0.1% of datapoints were lost. Cut offs were: max meal size 0.5g, meal duration 20 minutes and intermeal duration 80 mins.

Probability distribution functions (PDFs) for meal patterning from the CLAMS data were produced using customized MatLab scripts. We elected against a parametric assumption about the underlying (unknown) densities, and resorted to nonparametric kernel-based estimators. For n data on a given variable X, (x1,x2,…,xn) the kernel based estimate of the density of X at a generic point x is given by:

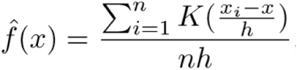

where K(·) is a kernel function that needs to satisfy certain properties and h is a bandwidth parameter that tends to zero as n tends to infinity. However, using a standard kernel function would yield inconsistent estimates near the origin since all the variables of interest in this dataset are non-negative by construction. In order to solve this potential problem, we used a simple reflection device^6^, together with the Epanechnikov kernel which has a bounded support. The bandwidth has been chosen using Silverman’s rule of thumb^7^. Statistical about the difference between comparator PDFS was via a Kolmogorov-Smirnov (K-S) test, which looks at the empirical distribution of a given variable and hence focuses on cumulative distribution functions rather than densities.

---

**Supplementary Figure 1.**
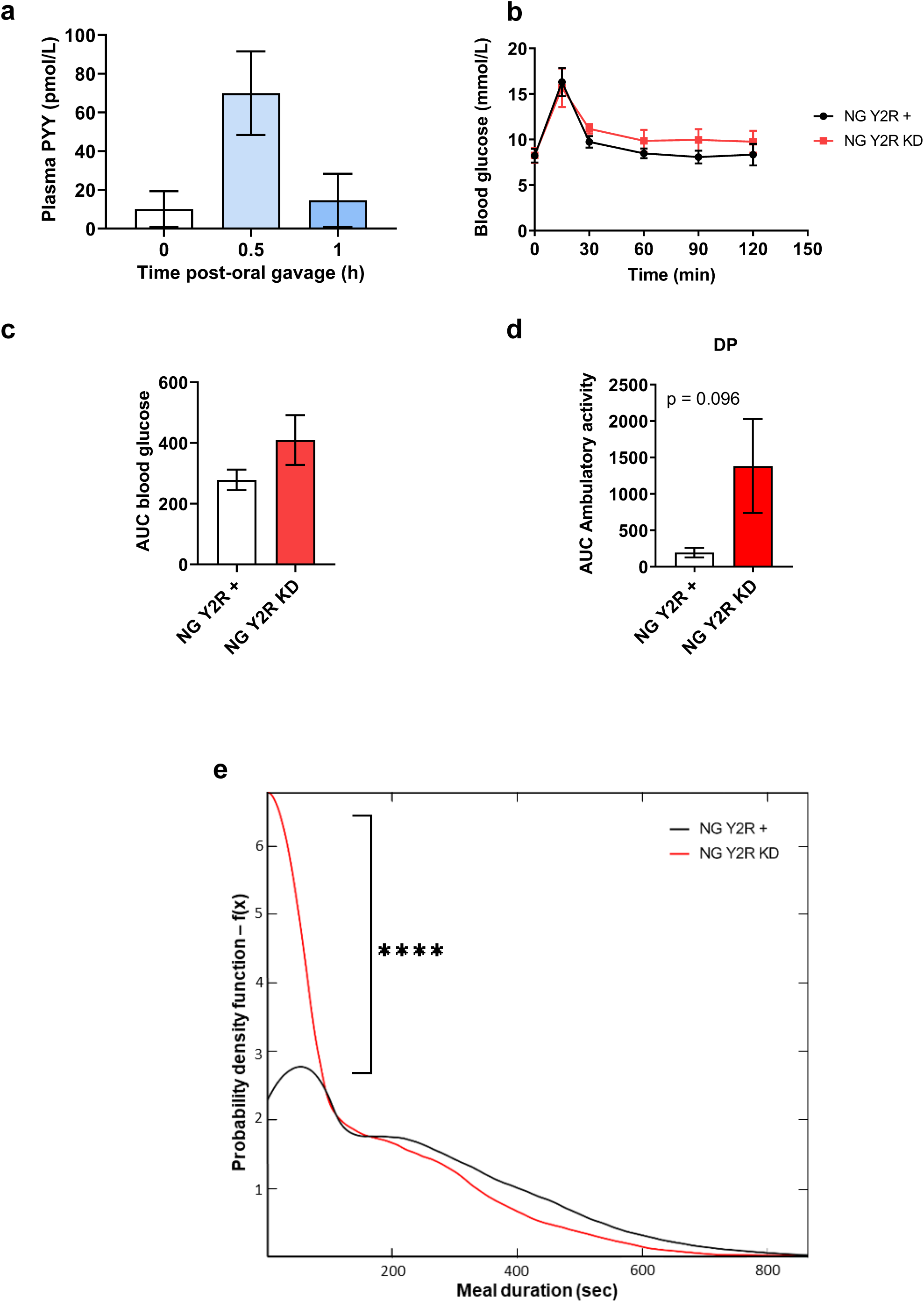
**S1a.** Plasma PYY levels in wild-type mice after oral gavage of the same meal bolus used in the feeding studies described in main Figure 2 (n = 4-5). **S1b and c**. Blood glucose in OGTT and its AUC during 0– 120 min in 4-h fasted mice (n = 7-5) in NG Y2R KD and control AAV-GFP NG injected animals (NG Y2R +). Two-way ANOVA followed by Dunnett’s test to compare to time 0 (in NG Y2R +, p < 0.01; in NG Y2R KD, p < 0.05). **S1d**. Area under the curve for ambulatory activity (beam breaks) in NG Y2R KD and AAV-GFP injected controls (mean of 3 days, dark phase in grey) in the CLAMS system (n = 6-6). Two-tailed unpaired t test. **S1e**. Probability density function of meal duration (i.e., time taken for feeding bout in seconds) over 7 days in a CLAMS system measuring high resolution food intake. The difference between NG Y2R KD vs NG Y2R + controls is highly significant (p<0.0001) Kolmogorov-Smirnov (K-S) test (n = 6 and 6).

